# ACTIVITY-DEPENDENT INTERNALIZATION OF GLUN2B-CONTAINING NMDARS IS REQUIRED FOR SYNAPTIC INCORPORATION OF GLUN2A AND SYNAPTIC PLASTICITY

**DOI:** 10.1101/2024.05.01.592099

**Authors:** Granville P. Storey, Andres Barria

**Author notes:** Corresponding author ORCID: 0000-0001-7724-0043.

## Abstract

NMDA-type glutamate receptors (NMDARs) are heterotetrameric complexes composed of two GluN1 and two GluN2 subunits. The precise composition of the GluN2 subunits determines the channel’s biophysical properties and influences its interaction with postsynaptic scaffolding proteins and signaling molecules involved in synaptic physiology and plasticity. Consequently, the precise regulation of NMDAR subunit composition at synapses is crucial for proper synaptogenesis, neuronal circuit development, and synaptic plasticity, a cellular model of memory formation.

In the forebrain during early development, NMDARs contain the GluN2B subunit, which is necessary for proper synaptogenesis and synaptic plasticity. In rodents, GluN2A subunit expression begins in the second postnatal week, replacing GluN2B-containing NMDARs at synapses in an activity- or sensory experience-dependent process. This switch in NMDAR subunit composition at synapses alters channel properties and reduces synaptic plasticity. The molecular mechanism regulating the switch remains unclear.

We have investigated the role of activity-dependent internalization of GluN2B-containing receptors in shaping synaptic NMDAR subunit composition. Using a combination of molecular, pharmacological, and electrophysiological approaches in cultured organotypic hippocampal slices from rats of both sexes, we show that the process of incorporating GluN2A-containing NMDARs receptors requires activity-dependent internalization of GluN2B-containing NMDARs. Interestingly, blockade of GluN2A synaptic incorporation was associated with impaired potentiation of AMPA-mediated synaptic transmission, suggesting a potential coupling between the trafficking of AMPARs into synapses and that of GluN2A-containing NMDARs.

These insights contribute to our understanding of the molecular mechanisms underlying synaptic trafficking of glutamate receptors and synaptic plasticity. They may also have implications for therapeutic strategies targeting NMDAR function in neurological disorders.

**SIGNIFICANCE STATEMENT:** Synaptic NMDARs play a critical role in synaptogenesis, synaptic stability, and activity-dependent regulation of synaptic strength. The developmental switch in GluN2 subunits composition of synaptic NMDARs is part of normal synapse development and is crucial for proper synaptic physiology, plasticity, and the formation of functional neuronal circuits, though the mechanisms governing it remain unclear.

We show that internalization of GluN2B-containing NMDARs is required for synaptic incorporation of GluN2A-containing receptors. This process can be induced by long-term potentiation and requires Ca^+2^. Notably, GluN2A trafficking to synapses is linked to the incorporation of AMPA-type glutamate receptors, suggesting a shared pathway for synaptic incorporation.

These findings provide greater insight into the molecular mechanisms behind glutamate receptor trafficking and synaptic plasticity, potentially informing therapeutic strategies for neurological disorders.

## INTRODUCTION

In many brain regions, synapses undergo a developmental switch in the subunit composition of NMDA receptors (Carmignoto and Vicini, 1992; Stocca and Vicini, 1998; Philpot et al., 2001; Yashiro and Philpot, 2008). During early development, NMDA receptors are composed of two of the obligatory GluN1 subunits, which is present in all NMDARs, and two GluN2B subunits, forming a tetramer (Hansen et al., 2021). GluN2B plays a crucial role in neural development, synaptic plasticity, and overall animal viability (Kutsuwada et al., 1996; Barria and Malinow, 2005; Ewald et al., 2008). It mediates prolonged channel openings, leading to long-lasting currents and subsequent prolonged postsynaptic depolarization. Additionally, its large intracellular carboxy-terminal domain allows the anchoring of active calcium/calmodulin-dependent kinase II (CaMKII) (Strack et al., 2000; Mayadevi et al., 2002), a necessary step for long-term potentiation of synaptic transmission mediated by AMPAR-type glutamate receptors (AMPARs) (Barria and Malinow, 2005; Zhou et al., 2007; Halt et al., 2012). Post-Golgi trafficking of GluN2B-containing NMDARs is regulated by palmitoylation (Hayashi et al., 2009; Mattison et al., 2012), while Wnt signaling can control the movement of GluN2B-containing NMDA receptors toward the neuronal surface (Cerpa et al., 2011; McQuate et al., 2017). From there, they can be incorporated into synapses by diffusion, independently of synaptic activity (Barria and Malinow, 2002; McQuate and Barria, 2020). GluN2B can be internalized in an activity-dependent manner through a YEKL motif in its intracellular carboxy-terminal domain, which interacts with the AP-2 complex (Lavezzari et al., 2003; Lavezzari et al., 2004).

The expression of other GluN2 subunits is developmentally regulated and region-specific (Akazawa et al., 1994; Monyer et al., 1994). In the hippocampus, cortex, and other forebrain regions of rodents, GluN2A expression increases after the second postnatal week, leading to a partial replacement of GluN2B-containing NMDARs with those containing GluN2A at synapses (Carmignoto and Vicini, 1992; Stocca and Vicini, 1998; Yashiro and Philpot, 2008; Storey et al., 2011). Consequently, mature synaptic NMDARs exhibit a range of compositions, including diheteromeric GluN1/GluN2B, diheteromeric GluN1/GluN2A, as well as triheteromeric GluN1/GluN2B/GluN2A receptors, with their ratios varying across different synapses (Hansen et al., 2021).

The switch in subunit composition of synaptic NMDARs alters the NMDAR-mediated current, as receptors containing GluN2A desensitize faster and exhibit faster opening and closing kinetics compared to those containing GluN2B (Dingledine et al., 1999). It also reduces synaptic plasticity, as the carboxy-terminal domain of GluN2A lacks a binding sequence for CaMKII (Omkumar et al., 1996; Mayadevi et al., 2002; Barria and Malinow, 2005). Different rules seem to govern trafficking of GluN2A-containing receptors into synapses, as they require synaptic activity acting on preexisting GluN2B-containing NMDARs for synaptic incorporation (Barria and Malinow, 2002). This process aligns with the observation that GluN2A-containing receptors are predominantly localized to synaptic sites (Groc et al., 2006; Kellermayer et al., 2018). Sensory experience is also required for the proper switch in the GluN2 subunit composition of synaptic NMDARs, with dark-reared animals maintaining a high GluN2B/GluN2A ratio at synapses in the visual cortex (Quinlan et al., 1999a; Quinlan et al., 1999b; Philpot et al., 2001). The carboxy-terminal domain of GluN2A lacks the YEKL motif, which mediates receptor endocytosis in GluN2B, thereby contributing to the greater stability of GluN2A-containing receptors at synapses (Lavezzari et al., 2003; Lavezzari et al., 2004; Sanz-Clemente et al., 2010).

Synaptic incorporation of GluN2A-containing NMDA receptors is typically detected by a decrease in the decay time of evoked excitatory postsynaptic currents (EPSCs). This reduction is commonly observed during the second postnatal week in the hippocampus of rodents—a period characterized by robust synaptogenesis and plasticity (Carmignoto and Vicini, 1992; Stocca and Vicini, 1998; Philpot et al., 2001; Storey et al., 2011). Dysregulation of the switch between GluN2B and GluN2A subunits could significantly impact synaptic plasticity and the development of neuronal properties (Gambrill and Barria, 2011), ultimately affecting the formation of functional brain circuits. In a mouse model of Fragile X syndrome, knockout of the FMR1 gene results in early expression and synaptic incorporation of GluN2A subunits (Banke et al., 2024). This alters the synaptic physiology and plasticity of neurons occurring during a critical period when synapses and circuits are developing, suggesting that dysregulation of the GluN2B/GluN2A switch may contribute to the formation of pathological circuits in the Fragile X mouse model.

The process through which the synaptic switch in GluN2 subunits is coordinated remains largely unknown. In this study, we specifically investigated whether the synaptic activity-induced removal of GluN2B is a prerequisite for the incorporation of GluN2A-containing receptors into synapses and examined the specific requirements for GluN2A synaptic incorporation.

## MATERIALS AND METHODS

### Slice cultures and transfection

Organotypic hippocampal slices were prepared from p6 Sprague-Dawley rats of both sexes as described previously (Opitz-Araya and Barria, 2011) and cultured at 35°C and 5% CO2 in medium containing 8.4 g/l MEM, 20% Horse serum heat inactivated, 1 mM L-Glutamine, 1 mM CaCl2, 2 mM MgSO4, 1 mg/l Insulin, 0.00125% Ascorbic Acid, 13 mM D-Glucose, 5.2 mM NaHCO3, and 30 mM and Hepes.

After 3-5 days in culture, slices were transfected using a biolistic particle delivery system (Woods and Zito, 2008) and cultured for an additional 48-72 hours to allow expression of recombinant proteins. For experiments requiring treatment of the slices with APV (Tocris), the drug was added immediately after transfection and replenished every 24 hours. In experiments requiring treatment with the antennapedia fusion peptides, 5 μM of the peptide was added to the slice culture media 2 hours before recording. In experiments requiring treatment with the casein kinase II inhibitor, TBB (Tocris), 10 μM was added to the slice culture media 2 hours before recording. In the case of TBB, 10 μM was also added to the bath solution during recording.

### Electrophysiology

Whole-cell recordings were obtained from transfected or non-transfected neurons under visual guidance using epi-fluorescence and transmitted-light illumination from an Olympus BX51W1 microscope. The recording chamber was perfused with artificial cerebrospinal fluid (ACSF) containing 119 mM NaCl, 2.5 mM KCl, 4 mM CaCl2, 4 mM MgCl2, 26 mM NaHCO3, 1 mM NaH2PO4, 11 mM glucose, 100 μM picrotoxin (Tocris), 2 μM 2-chloroadenosine (pH 7.4), and gassed with 5% CO2 / 95% O2. Recordings were performed at room temperature (20-25°C), except in rapid incorporation experiments which were performed at 27°C. Whole-cell recording pipettes (3-6 MΩ) were filled with intracellular recording solution containing 115 mM cesium methanesulfonate, 20 mM CsCl, 10 mM HEPES, 2.5 mM MgCl2, 2 mM MgATP, 2 mM Na2ATP, 0.4 mM Na3GTP, 10 mM sodium phosphocreatine, 5 mM QX-314, and 0.6 mM EGTA (pH 7.25 and 310 mmol/Kg. Whole-cell recordings were made using a Multiclamp 700B computer-controlled amplifier (Axon Instruments). Synaptic responses were evoked by bipolar cluster electrodes (FHC) placed over Schaffer Collateral fibers approximately 200-300 μm from targeted CA1 neuron. NMDAR kinetics were measured as time to half-decay (100% peak current to 50% remaining current) obtained from evoked EPSCs while holding the cell +40 mV in the presence of 2 μM NBQX. In experiments measuring incorporation index of recombinant GluN2A, the NMDAR amplitude was measured as the mean current from a 50 ms window 110 ms after the stimulus artifact in evoked EPSCs recorded while holding the cell at -60 mV. The peak amplitude of the EPSC is mediated by endogenous AMPARs and was used to normalize the recombinant NMDAR response.

### Plasmids and Peptide construction

A mammalian expression plasmid carrying the YEKL domain of GluN2B was constructed by fusing mCherry to the GluN2B carboxy-terminal at A1315 and incorporating a stop codon after L1475, thus eliminating the PDZ binding motif. A control plasmid was created by inserting a stop codon after V1471, thus also eliminating the YEKL motif.

GluN2A and GluN1 N598R were tagged with GFP and expressed using a mammalian expression plasmid as reported previously (Barria and Malinow, 2002).

Antennapedia peptides were synthesized by Genemed Synthesis containing the following amino acid sequences: RQIKIWFQNRRMWKKNGHVYEKLSSIE for the Ant-YEKL peptide and RQIKIWFQNRRMKWKKNGHVAEKLSSIE for the control Ant-AEKL peptide.

### Experimental Design and Statistical Analysis

Whole-cell recordings were obtained from one or two neurons per hippocampal slice. Batches of organotypic cultured slices were obtained from 3 to 6 Sprague-Dawley rats of both sexes at postnatal day 6 (P6) and cultured for 6-8 days. All results are presented as mean ± SEM.

Statistical comparisons between paired recordings (Figures 1 and supplemental) were conducted using a non-parametric Wilcoxon test, while comparisons between control and treated conditions (Figures 4 and 5) used an unpaired non-parametric Mann-Whitney test. A two-tailed p-value was applied, with the specific p-values indicated in each figure legend. Statistical significance was defined as p < 0.05.

**Figure 1.**
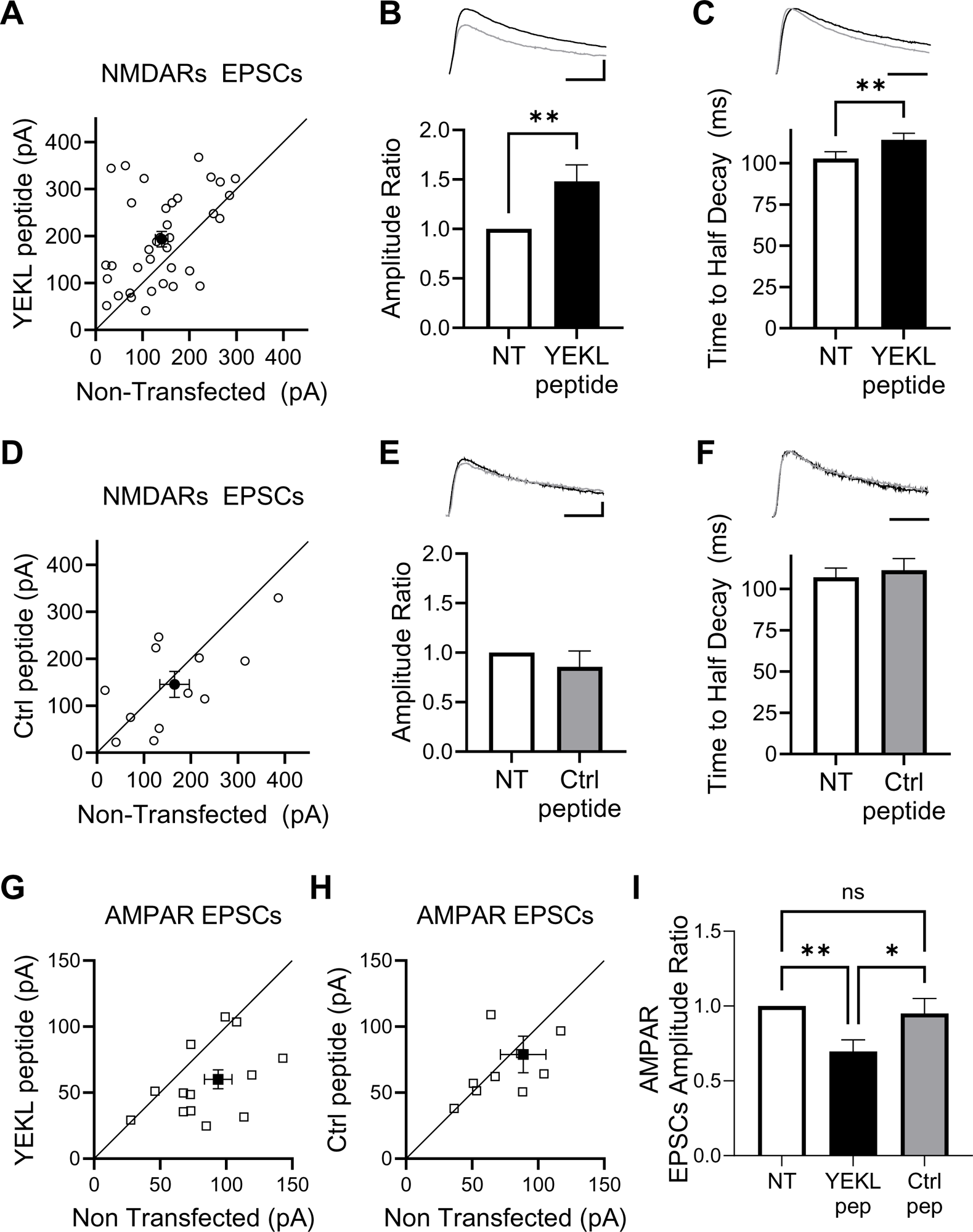
Blockade of endogenous GluN2B-containing receptors internalization by YEKL peptide. **A.** Paired recordings of neurons transfected with YEKL peptide and control non-transfected (NT) neurons. Peak amplitude of isolated NMDAR-mediated currents recorded at +40mV and in the presence of 2 mM NBQX from CA1 transfected and control neurons recorded as pairs (n=36 pairs). Optically tagged YEKL peptide was transfected using biolistic and expressed for 2-3 days. Black dot is average ± s.e.m. **B.** Top, sample EPSC traces from a control neuron (gray trace) and a neuron expressing YEKL peptide (black trace) recorded at +40 mV. Bars = 50 ms and 100 pA. Bottom, peak amplitude of NMDAR-mediated EPSCs from paired recordings normalized to the amplitude of the control non-transfected (NT) neuron. Statistical significance was determined using Wilcoxon matched pairs test (p=0.0049). **C.** Top, sample EPSC traces normalized to peak amplitude from a control neuron (gray trace) and a neuron expressing YEKL peptide (black trace) recorded at +40 mV. Bar = 50 ms. Bottom, Time to Half Decay of NMDAR EPSCs from control non-transfected (NT) neurons and neurons expressing YEKL peptide. Statistical significance was determined using Wilcoxon matched pairs test (p=0.0012). **D.** Paired recordings of neurons transfected with control peptide lacking the YEKL motif and control non-transfected (NT) neurons. Peak amplitude of isolated NMDAR-mediated currents recorded at +40mV and in the presence of 2 mM NBQX from CA1 transfected and control neurons recorded as pairs (n=12 pairs). Control peptide was transfected using biolistic and expressed for 2-3 days. Black dot is average ± s.e.m. **E.** Top, sample EPSC traces from a control neuron (gray trace) and a neuron expressing control peptide (black trace). Bar = 50 ms and 50 pA. Bottom, peak amplitude of NMDAR-mediated EPSCs from paired recordings normalized to the amplitude of the control non-transfected (NT) neuron. Statistical significance was determined using Wilcoxon matched pairs test (p=0.3203). **F.** Top, sample EPSC traces normalized to peak amplitude from a control neuron (gray trace) and a neuron expressing control peptide (black trace). Bar = 50 ms. Bottom, Time to Half Decay of NMDAR EPSCs from control non-transfected (NT) neurons and neurons expressing control peptide. Statistical significance was determined using Wilcoxon matched pairs test (p=0.5693). **G-H.** Peak amplitude of paired recordings of AMPAR-mediated EPSCs recorded at -60 mV from CA1 neurons expressing either YEKL peptide (G, n=15 pairs) or the control peptide (H, n=10 pairs), and an adjacent non-transfected cell (NT). Black square is average ± s.e.m. **I.** Peak amplitude of AMPAR-mediated EPSCs from paired recordings of transfected neurons as indicated normalized to the amplitude of the control non-transfected (NT) neuron. Statistical significance was determined using one-way ANOVA (F=6.578; p=0.0036) followed by Tukey’s multiple comparison test (*** p=0.0009; * p=0.0363)

For comparisons involving multiple treatments (Figures 2 and 3), a one-way ANOVA was performed, followed by Tukey’s post hoc test. The F and p values for each ANOVA are listed in the figure legends, along with the specific p values from Tukey’s post hoc test. All data were analyzed using Prism software.

**Figure 2.**
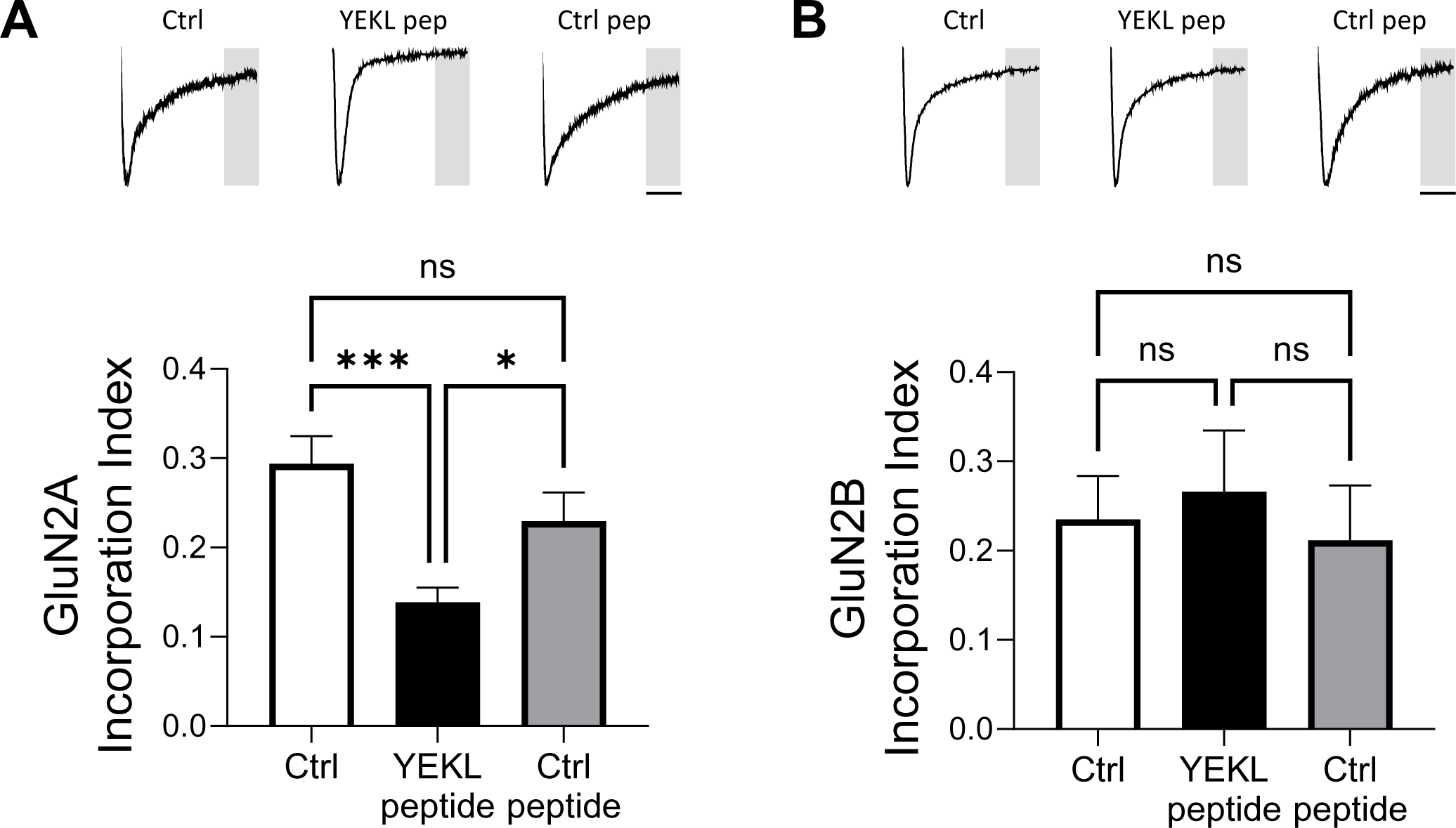
Synaptic incorporation of recombinant GluN2A-containing receptors. **A.** Top, sample EPSCs recorded at -70 mV from CA1 neurons co-transfected with GFP-tagged NMDAR subunits GluN2A and GluN1 N598R, and either YEKL peptide or control peptide. Control cells (far left trace) were co-transfected only with GFP-tagged NMDAR subunits GluN2A and GluN1 N598R. Bar=50 ms. Bottom, GluN2A Incorporation Index from cells as indicated measured as the average current 150 msec after the stimulus artifact (grey area) normalized to the peak amplitude occurring shortly after the stimulus artifact. Statistical significance was determined using one-way ANOVA (F=7.812; p=0.0010) followed by Tukey’s multiple comparison test (** p=0.0041; * p=0.0364). Control cells n=15, cells co-expressing YEKL peptide and control peptide n=23 each. **B.** Top, sample EPSCs recorded at -70 mV from CA1 neurons co-transfected with GFP-tagged NMDAR subunits GluN2B and GluN1 N598R, and either YEKL peptide or control peptide. Control cells (far left trace) were co-transfected only with GFP-tagged NMDAR subunits GluN2B and GluN1 N598R. Bar=50 ms. Bottom, GluN2B Incorporation Index from cells as indicated and measured as in A. Statistical significance was determined using one-way ANOVA (F=0.1355; p=0.8737). Control cells n=14, cells co-expressing YEKL peptide n=16, and cells co-expressing control peptide n=5.

**Figure 3.**
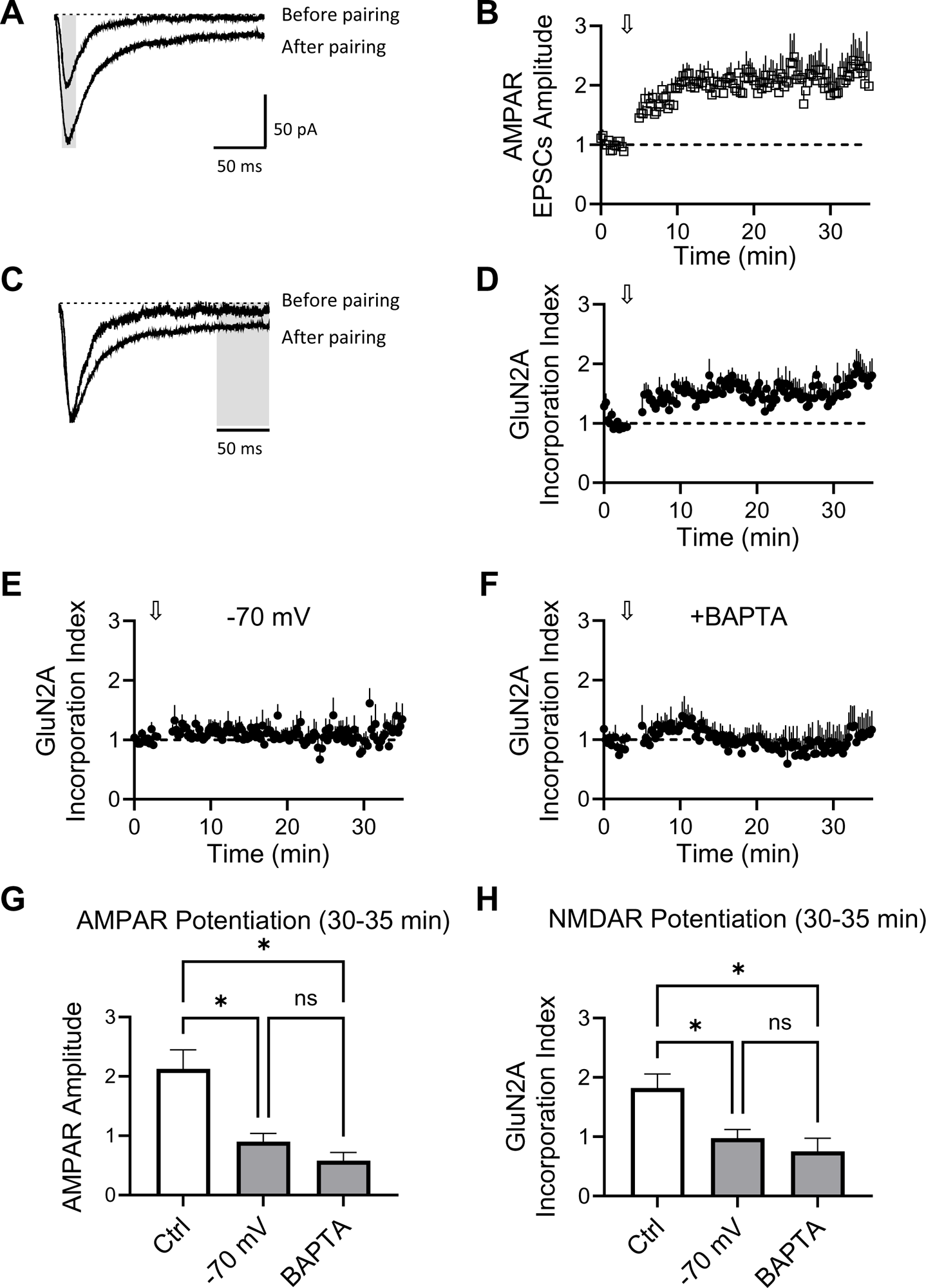
Rapid synaptic incorporation of GluN2A-containing receptors. **A.** Sample traces of evoked excitatory postsynaptic currents (EPSCs) recorded at -70 mV from a CA1 neuron co-transfected with GFP-tagged NMDAR subunits GluN2A and GluN1 N598R. The traces show responses during baseline (before a pairing protocol) and 20 minutes after potentiation, which was induced by a 2-minute, 3 Hz stimulation while holding the cell at 0 mV. **B.** Peak amplitude of EPSCs from CA1 neurons as in Figure A, with the gray area indicating the AMPAR-mediated component of the EPSCs. The arrow indicates the point where the pairing protocol was applied to induce potentiation (n=15). **C.** Sample traces as in A normalized to the peak amplitude of the response. The recombinant NMDAR current is quantified 150-200 msec after the stimulus artifact (gray area). **D.** Amplitude of the current carried by GluN2A-containing recombinant NMDARs over time. The arrow indicates the application of the pairing protocol, which involves 2 minutes of 3 Hz stimulation while holding the cell at 0 mV. **E.** Amplitude of the current carried by GluN2A-containing recombinant NMDARs in experiments with a pairing protocol consisting of 2 minutes of 3 Hz stimulation while holding the cell at -70 mV (n=23). **F.** Amplitude of the current carried by GluN2A-containing recombinant NMDARs over time from CA1 neurons recorded with internal solution containing BAPTA. The arrow indicates the application of the pairing protocol, which involves 2 minutes of 3 Hz stimulation while holding the cell at 0 mV (n=12). **G-H.** Quantification of potentiation 30-35 minutes after the start of the experiment, measuring the AMPAR-mediated component of the EPSC (G) and the late component of the EPSC resulting from the synaptic expression of GluN2A-containing recombinant NMDARs (H). Statistical significance was determined using one-way ANOVA (AMPAR potentiation F=6.645 and p=0.0058; NMDAR potentiation F=6.452 and p=0.0065) followed by Tukey’s multiple comparison test. Asterisk p<0.05.

## RESULTS

### Blockade of activity-dependent internalization of endogenous GluN2B containing receptors

Activity-dependent internalization of GluN2B containing NMDARs requires the interaction of the endocytic motif YEKL present only in the C-terminus of the GluN2B subunit, and the clathrin adaptor protein AP-2. Phosphorylation of tyrosine 1472 within the endocytic motif by Src kinases prevents this interaction, thereby stabilizing the surface expression of NMDARs (Roche et al., 2001; Lavezzari et al., 2003; Zhang et al., 2008; Sanz-Clemente et al., 2010).

We aimed to disrupt the internalization of GluN2B-containing NMDARs in CA1 pyramidal neurons by expressing a competing peptide containing the YEKL motif. We hypothesized that this peptide would competitively inhibit the interaction between endogenous GluN2B and the AP-2 endocytic machinery, thereby blocking activity-dependent GluN2B internalization and enhancing its presence at synapses.

Organotypic cultured hippocampal slices (Opitz-Araya and Barria, 2011) were transfected using biolistic (Woods and Zito, 2008) with an mCherry-tagged peptide named YEKL peptide. This peptide contained the last portion of the GluN2B C-tail (amino acids 1315-1475) but lacked the PDZ binding domain to prevent interactions with proteins from the MAGUK family (Prybylowski et al., 2005). As a control, a similar construct lacking the YEKL motif (amino acids 1315-1471) was utilized. Following 36-60 hours of expression, we assessed the impact of these constructs on endogenous synaptic NMDARs by comparing the amplitude of isolated NMDAR-evoked postsynaptic currents (EPSCs) recorded from a transfected neuron and an adjacent non-transfected neuron (paired recordings). EPSCs were evoked by stimulating Schaffer collaterals with bipolar electrodes.

Cells expressing the YEKL peptide exhibited larger (Figure 1A and 1B) and slower (Figure 1C) NMDA-R EPSCs compared to adjacent non-transfected cells stimulated under the same conditions. The increase in the time to half decay is minimal, as expected for an EPSC already dominated by GluN2B; however, it is consistent with an increased GluN2B to GluN2A ratio at synapses. Neurons transfected with the control peptide showed no increase in amplitude or change in kinetics of the NMDAR-mediated EPSCs compared to adjacent non-transfected cells (Figure 1D-F).

We also investigated the impact of expressing the YEKL peptide on synaptic AMPA-type glutamate receptors (AMPARs). Surprisingly, blocking the internalization of GluN2B-containing NMDARs led to a decrease in the amplitude of AMPAR-mediated EPSCs compared to adjacent non-transfected neurons (Figure 1G and 1I). This observation could be attributed to the disruption of normal synaptic plasticity. Specifically, the forward trafficking of AMPARs and their synaptic incorporation, which typically occurs in cultured organotypic slices during the expression period of the peptides, may be affected. The expression of the control peptide had no effect on AMPAR-mediated synaptic transmission (Figure 1H and 1I).

To assess whether the YEKL peptide affected the removal or recycling of other membrane receptors, we examined the amplitude of isolated GABA receptor IPSCs. We chose to investigate GABA receptor transmission because GABA-R endocytosis is known to be regulated by AP-2 (Kittler et al., 2000; Kneussel, 2002). Recording from pairs consisting of a YEKL-transfected neuron and a non-transfected neighbor, we evoked IPSCs in the presence of 100 μM APV and 2 μM NBQX and compared amplitudes. However, we did not observe any differences in the amplitude of the GABA-R IPSCs, leading us to conclude that the removal or recycling of GABA-Rs was not affected by YEKL peptide expression (Supplemental Figure 1A-B).

This data indicates that the YEKL peptide competes with the tail of endogenous GluN2B for binding of AP-2, thereby preventing the internalization of synaptic GluN2B receptors and leading to an increase in the number of synaptic GluN2B-containing NMDARs relative to non-transfected paired cells, consistent with previous reports (Lavezzari et al., 2003; Lavezzari et al., 2004). These findings validate the use of the YEKL peptide to interfere with NMDAR internalization. Furthermore, these experiments suggest that the internalization of GluN2B-containing receptors may be coupled to the normal synaptic incorporation of AMPA-type glutamate receptors during the expression period in the cultured slice.

### YEKL peptide blocks synaptic incorporation of recombinant GluN2A-containing receptors

In rodents, expression of the GluN2A subunits starts around postnatal day 10 (p10) (Storey et al., 2011) and immediately is incorporated into synapses in a process that requires synaptic or sensory activity (Quinlan et al., 1999a; Barria and Malinow, 2002) and it is also induced during synaptic potentiation using well established long-term potentiation (LTP) protocols (Bellone and Nicoll, 2007).

We hypothesize that the synaptic incorporation of GluN2A-containing NMDARs is dependent on the prior removal of GluN2B-containing receptors. To test this, we used an electrophysiologically tagged GluN2A-containing NMDARs that allowed us to functionally quantify its incorporation into synapses (Barria and Malinow, 2002).

In brief, GluN2A was co-expressed with a mutant form of GluN1 (GluN1 N598R) that eliminates the normal Mg^2+^ blockade of NMDARs observed at resting membrane potentials (Burnashev et al., 1992; Single et al., 2000). Both subunits were optically tagged with GFP to identify transfected CA1 neurons in the slice. In transfected cells, EPSCs evoked at -60 mV exhibit a fast component due to the activation of endogenous AMPARs and a slow component that reflects the activation of recombinant NMDARs, as previously described (Barria and Malinow, 2002). A synaptic incorporation index was calculated by measuring the late component of the EPSC, which occurs 150-200 ms after the stimulus artifact and reflects the synaptic activation of recombinant NMDARs (Figure 2, gray area on sample traces). This measurement was then normalized to the peak amplitude of the fast component, which reflects the activation of endogenous AMPARs.

We recorded excitatory postsynaptic currents (EPSCs) at -60 mV in control cells transfected with electrophysiologically tagged GluN2A receptors. As illustrated in Figure 2A, a late current indicative of the synaptic incorporation of recombinant GluN2A-containing NMDARs is observed. However, in cells co-expressing the YEKL peptide, which blocks synaptic removal of GluN2B, this current was absent, suggesting that GluN2A-containing NMDARs were not incorporated into synapses. In contrast, GluN2A-containing NMDARs were normally incorporated into synapses in cells co-expressing the control peptide (Fig 2A).

We also measured the synaptic incorporation of electrophysiologically tagged GluN2B-containing receptors in a similar manner. We found that the YEKL peptide had no effect on the incorporation of recombinant GluN2B-containing receptors, consistent with previous reports indicating that forward trafficking or synaptic incorporation of GluN2B is independent of synaptic activity (Fig 2B) (Barria and Malinow, 2002).

These experiments suggest that the internalization of GluN2B, likely induced by the normal synaptic activity that occurs in the cultured organotypic slices during the expression period, is required for the synaptic incorporation of recombinant GluN2A.

### Requirements for synaptic incorporation of recombinant GluN2A-containing receptors

To better understand the synaptic incorporation of GluN2A-containing NMDARs in response to a more defined pattern of synaptic activity, we once again utilized electrophysiologically tagged GluN2A-containing receptors to monitor their synaptic incorporation in real-time.

Organotypic hippocampal slices, cultured for 3-5 days, were co-transfected with GluN1 N598R and GluN2A. Subsequently, the slices were cultured for an additional 3 days in the presence of 100 μM APV. This treatment was employed to allow for the expression of the receptor without its synaptic incorporation, as synaptic activity is required for the incorporation of GluN2A into synapses (Barria and Malinow, 2002; Storey et al., 2011).

We recorded evoked EPSCs from CA1 pyramidal cells expressing recombinant NMDARs while holding the membrane potential at -70 mV. Schaffer collaterals were stimulated at a frequency of 0.3 Hz, and the amplitude of the fast component reflecting AMPAR-mediated current was recorded (Figure 3A and 3B). Subsequently, synaptic potentiation was induced using a pairing protocol consisting of 3 Hz stimulation for 2 minutes, with the membrane potential held at 0 mV (Figure 3B, arrow). Figure 3B quantifies the increase in amplitude of AMPAR-mediated responses, a well-documented synaptic potentiation of AMPAR-mediated transmission following this pairing protocol (Barria and Malinow, 2005).

To estimate synaptic incorporation of recombinant GluN2A-containing receptors, we measured the late component at 150-200 ms and normalized that value to the peak amplitude of the fast component (Figure 3C, grey area). Before pairing, no recombinant GluN2A-mediated current is observed as expected because APV had blocked the incorporation of recombinant GluN2A (Figure 3C, before pairing). Following the pairing protocol, a rapid increase in this current is clearly observed, indicating rapid incorporation of recombinant GluN2A-containing receptors (Figure 3D).

These experiments establish that a widely used LTP inducing protocol, in addition to augmenting AMPAR-mediated synaptic transmission, also drives synaptic incorporation of GluN2A-containing receptors.

Next, we employed this approach to examine whether synaptic activity alone, specifically glutamate release from presynaptic terminals, was adequate to acutely drive the synaptic incorporation of recombinant GluN2A, or if postsynaptic Ca^2+^ influx was necessary for this process.

In CA1 pyramidal cells expressing recombinant GluN2A-containing receptors, following a baseline period, we increased the stimulation rate to 3 Hz for 2 minutes while maintaining the cell at -70 mV to prevent the activation of endogenous NMDARs. Despite this brief period of increased activity, no significant changes in the late component of EPSCs were observed (Figure 3E), indicating that this level of activity was insufficient to drive the synaptic incorporation of GluN2A. This suggests that current flowing through existing synaptic NMDARs is necessary.

To confirm that calcium influx is indeed a requirement for GluN2A synaptic incorporation, we repeated the experiment with a calcium chelator in the pipette (10 mM BAPTA). The LTP-inducing protocol used previously failed to induce potentiation of the AMPA-R mediated synaptic transmission, as expected. Importantly, it also failed to induce synaptic incorporation of electrophysiologically tagged GluN2A-containing receptors (Figure 3F).

Figures 3G and 3H depict the quantification of the potentiation of AMPAR-mediated and recombinant NMDAR-mediated EPSCs, respectively, observed after 25-30 minutes of the pairing protocol under different conditions as indicated.

These experiments indicate that synaptic activity, through the activation of existing synaptic NMDARs and subsequent postsynaptic calcium influx, is necessary for GluN2A synaptic incorporation.

Taken together, these experiments suggest that the synaptic incorporation of GluN2A-containing receptors and the potentiation of AMPAR-mediated synaptic transmission are interconnected processes. Indeed, a positive correlation is observed when the potentiation of AMPA-R mediated synaptic transmission is plotted against the potentiation of the late component mediated by the newly inserted NR2A receptors (Supplemental Figure 1C).

### Removal of NR2B is required for rapid synaptic incorporation of NR2A

After directly examining the requirements for inducing the rapid incorporation of electrophysiologically tagged GluN2A-containing receptors into synapses, our next question was whether blockade of synaptic activity-induced GluN2B removal is necessary.

We coexpressed electrophysiologically tagged GluN2A receptors with either the YEKL peptide or the control peptide in organotypic slices. As previously, slices were maintained in 100 μM APV during the expression period (3 days in culture) to prevent the incorporation of GluN2A into synapses.

Figures 4A and 4B show the late component of evoked synaptic responses, which represents the amount of GluN2A that has incorporated into synapses. After 3-5 minutes of baseline recording, a pairing protocol of 3 Hz stimulation for 2 minutes with the cell voltage clamped at 0 mV was applied. Neurons co-expressing the control peptide exhibited a rapid incorporation of recombinant GluN2A-containing receptors, as shown in Figure 4A. In contrast, cells co-expressing the YEKL peptide did not show this incorporation, as illustrated in Figure 4B. This result further supports the notion that GluN2B internalization is necessary for the synaptic incorporation of GluN2A.

**Figure 4.**
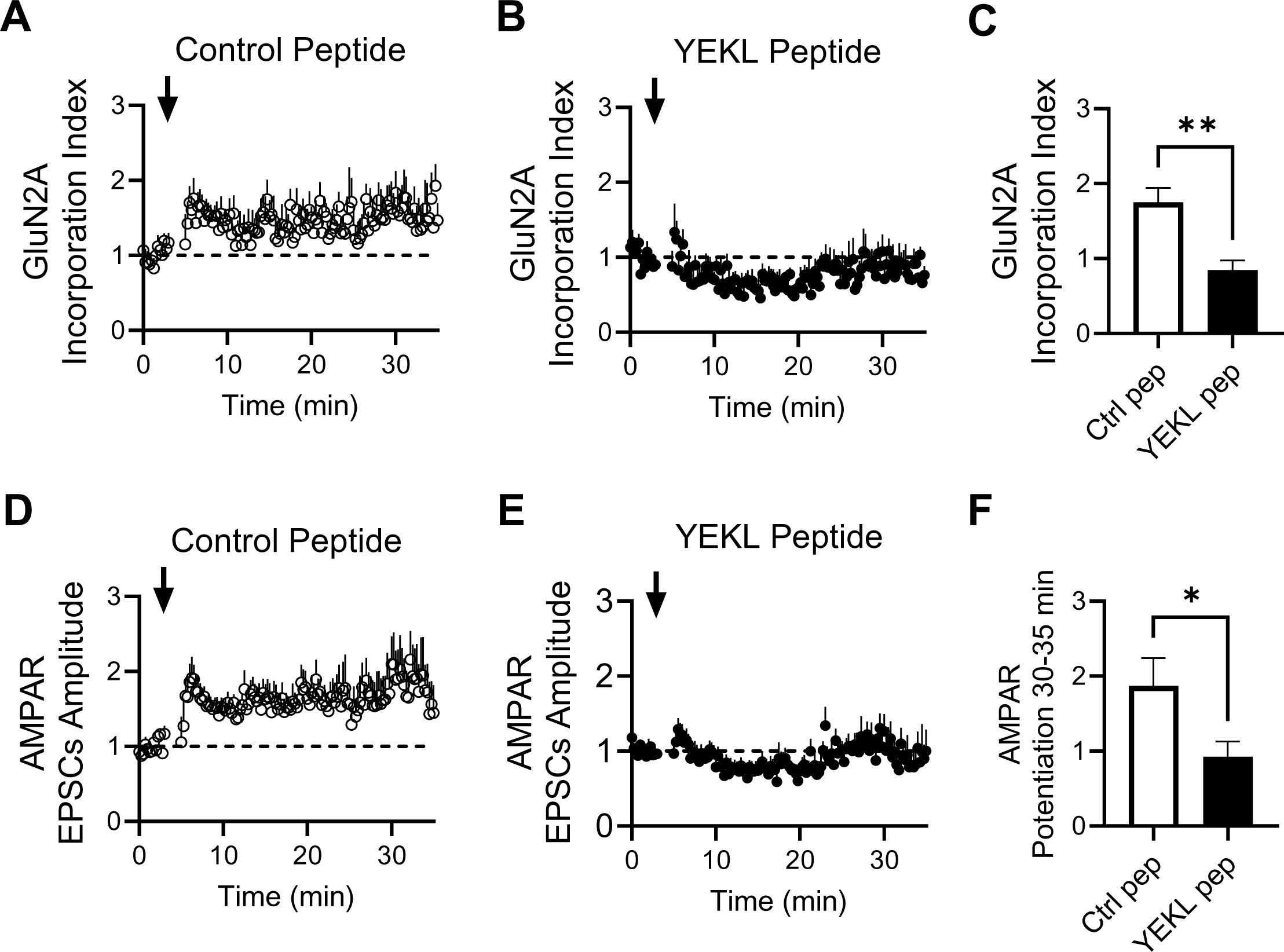
Synaptic incorporation of GluN2A-containing receptors while blocking internalization of GluN2B. **A.** Amplitude of the current carried by GluN2A-containing recombinant NMDARs over time in CA1 neurons co-transfected with GFP-tagged NMDAR subunits GluN2A and GluN1 N598R, along with the control peptide (n=14). EPSCs were recorded at -70 mV and the GluN2A current measured 150 ms after the stimulus artifact. The arrow indicates the induction of synaptic potentiation through a pairing protocol, which consists of 2-minute, 3 Hz stimulation while holding the cell at 0 mV. **B.** Amplitude of the current carried by GluN2A-containing recombinant NMDARs over time in CA1 neurons co-transfected with GFP-tagged NMDAR subunits GluN2A and GluN1 N598R, along with the YEKL peptide (n=10). EPSCs were recorded at -70 mV and the GluN2A current measured 150 ms after the stimulus artifact. The arrow indicates the induction of synaptic potentiation through a pairing protocol, which consists of 2-minute, 3 Hz stimulation while holding the cell at 0 mV. **C.** Quantification of GluN2A synaptic incorporation 30-35 min after the start of the experiment for neurons expressing either the control peptide or the YEKL peptide. Statistical significance was determined using Mann-Whitney test (p=0.0070). **D.** Peak amplitude of EPSCs recorded at -70 mV, representing the AMPAR component of the EPSC, in CA1 neurons as in A, i.e. expressing the control peptide. **E.** Peak amplitude of EPSCs recorded at -70 mV, representing the AMPAR component of the EPSC, in CA1 neurons as in B, i.e. expressing YEKL peptide. **F.** Quantification of AMPAR-mediated synaptic transmission 30-35 min after start of the experiment in neurons expressing either the control peptide or the YEKL peptide. Statistical significance was determined using Mann-Whitney test (p=0.0315).

Interestingly, expression of the YEKL peptide also prevented potentiation of AMPAR-mediated synaptic transmission (Figure 4E and F), adding to evidence suggesting that AMPA-R potentiation is linked to the incorporation of NR2A into the synapse. The control peptide had no effect on the synaptic potentiation of AMPAR (Figure 4D and F).

### Acute Blockade on GluN2B internalization also prevents GluN2A synaptic incorporation and AMPAR potentiation

We were concerned that prolonged blockade of GluN2B removal via expression of the YEKL peptide could potentially lead to a misinterpretation of our results. To address this concern, we next examined the effects of acute blockade of GluN2B removal on the rapid synaptic incorporation of GluN2A and AMPAR potentiation.

We used an antennapedia fusion peptide containing the YEKL domain of the GluN2B carboxy-tail (Ant-YEKL). The antennapedia sequence renders our YEKL domain peptide membrane-permeable, allowing for its brief application directly to the slice culture media before recordings (Fink et al., 2003; Illario et al., 2003; Schmitt et al., 2004; Sanhueza et al., 2007). Additionally, we developed a control antennapedia fusion peptide named Ant-AEKL, which incorporated the same GluN2B C-tail sequence as Ant-YEKL but with the tyrosine (Y) replaced by an alanine (A), thus preventing this peptide from binding to AP-2 (Prybylowski et al., 2005).

Cultured organotypic hippocampal slices were transfected with electrophysiologically tagged GluN2A-containing receptors and cultured with 100μM APV to prevent synaptic incorporation during the expression period due to spontaneous activity in the slice (Barria and Malinow, 2002).

The slices were treated with either 5μM of Ant-YEKL or Ant-AEKL for two hours before recording evoked currents from transfected CA1 pyramidal neurons, while maintaining the cell at -70 mV. As before, we measured the fast component of the evoked EPSCs, indicative of AMPA-mediated transmission, and the late component of the EPSC, indicative of electrophysiologically tagged GluN2A presence at synapses.

After establishing a stable baseline for 3 to 5 minutes, we applied a pairing protocol consisting of 3 Hz stimulation while holding the cell at a voltage of 0 mV.

In slices treated with the control Ant-AEKL peptide, we observed rapid synaptic incorporation of GluN2A-containing receptors (Figure 5A), along with the expected potentiation of AMPAR-mediated EPSCs (Figure 5D).

**Figure 5.**
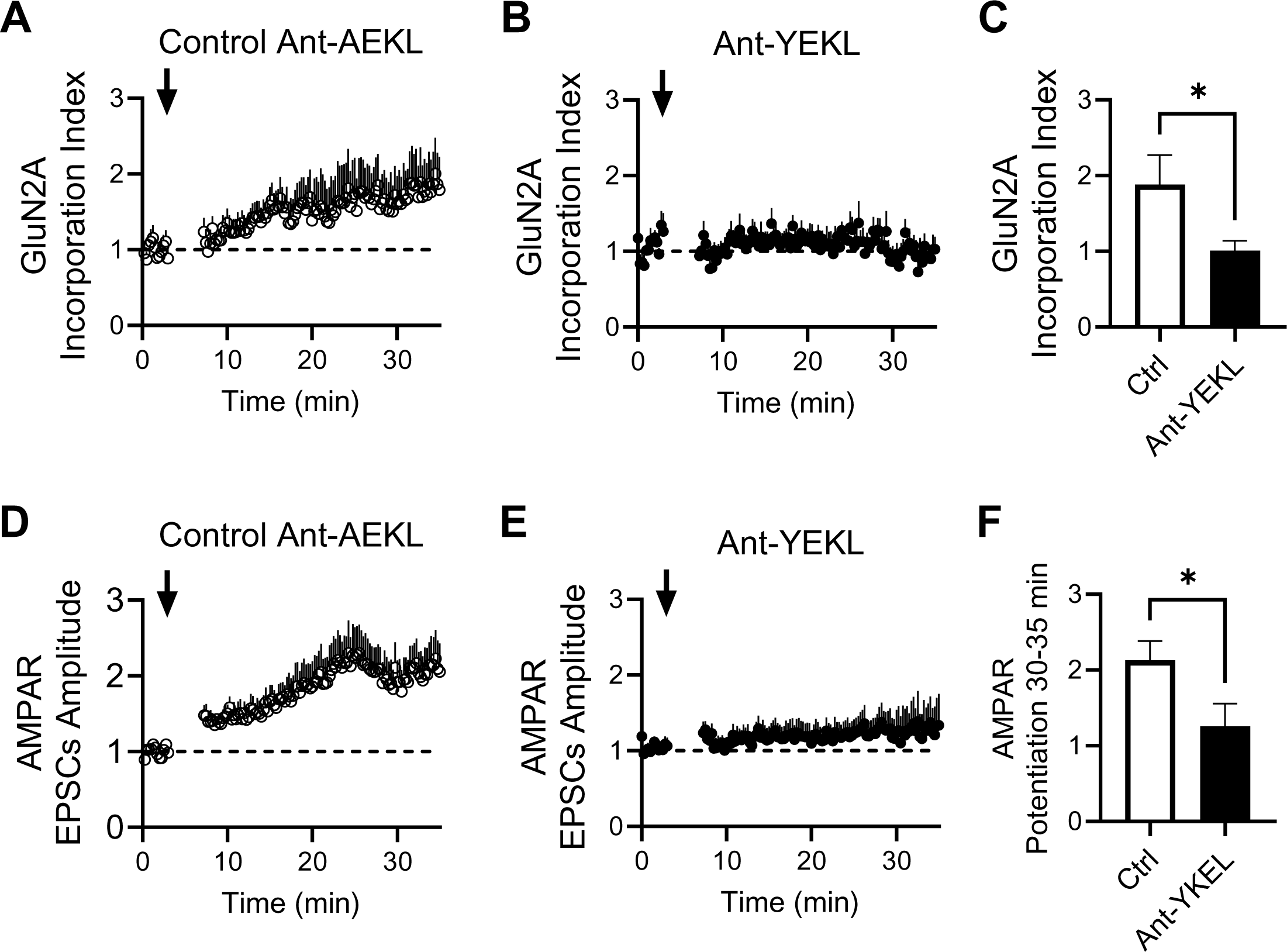
Rapid blockade of GluN2B internalization and its effect on GluN2A synaptic incorporation. **A.** Amplitude of the current carried by GluN2A-containing recombinant NMDARs over time in CA1 neurons co-transfected with GFP-tagged NMDAR subunits GluN2A and GluN1 N598R. Slices were incubated with the control antennapedia peptide Ant-AEKL for 2 hours before the experiment (n=9). EPSCs were recorded at -70 mV and the GluN2A current measured 150 ms after the stimulus artifact. The arrow indicates the induction of synaptic potentiation through a pairing protocol, which consists of 2-minute, 3 Hz stimulation while holding the cell at 0 mV. **B.** Amplitude of the current carried by GluN2A-containing recombinant NMDARs over time in CA1 neurons co-transfected with GFP-tagged NMDAR subunits GluN2A and GluN1 N598R. Slices were incubated with the antennapedia peptide Ant-YEKL for 2 hours before the experiment (n=9). EPSCs were recorded at -70 mV and the GluN2A current measured 150 ms after the stimulus artifact. The arrow indicates the induction of synaptic potentiation through a pairing protocol, which consists of 2-minute, 3 Hz stimulation while holding the cell at 0 mV. **C.** Quantification of GluN2A synaptic incorporation 30-35 minutes after the start of the experiment for neurons treated with either the control Ant-AEKL peptide or the Ant-YEKL peptide. Statistical significance was determined using Mann-Whitney test (p = 0.0293). **D.** Peak amplitude of EPSCs recorded at -70 mV, representing the AMPAR component of the EPSC, in CA1 neurons coexpressing GFP-tagged NMDAR subunits GluN2A and GluN1 N598R and treated with control antennapedia peptide Ant-AEKL. **E.** Peak amplitude of EPSCs recorded at -70 mV, representing the AMPAR component of the EPSC, in CA1 neurons coexpressing GFP-tagged NMDAR subunits GluN2A and GluN1 N598R and treated with antennapedia peptide Ant-YEKL. **F.** Quantification of AMPAR-mediated synaptic transmission 30-35 minutes after the start of the experiment for neurons treated with either the control Ant-AEKL peptide or the Ant-YEKL peptide. Statistical significance was determined using Mann-Whitney test (p = 0.0418).

In contrast, in slices treated with the Ant-YEKL peptide synaptic incorporation of GluN2A-containing receptors was prevented (Figure 5B and C) as well as potentiation of AMPAR-mediated currents (Figure 5E and F)

We also used a pharmacological approach to block activity-dependent internalization of GluN2B-containing receptors. We used the casein kinase II inhibitor, TBB. Phosphorylation of GluN2B by casein kinase II has been shown to be a critical step in triggering activity-dependent removal of GluN2B from the synapse (Sanz-Clemente et al., 2010).

Hippocampal slices expressing electrophysiologically tagged GluN2A receptors were treated with 10μM TBB for two hours before recording. As before, slices were cultured with 100 μM APV to prevent incorporation of GluN2A during the expression period.

In Figure 6A, consistent with the outcomes observed with the expression of the YEKL peptide (Figures 2 and 4) and treatment with Ant-YEKL peptide (Figure 5), pharmacological inhibition of GluN2B synaptic removal impeded the rapid incorporation of GluN2A induced by the pairing protocol. Moreover, the prevention of GluN2A incorporation coincided with the absence of potentiation in the AMPAR-mediated component of the EPSC, aligning with the findings from other experiments.

**Figure 6.**
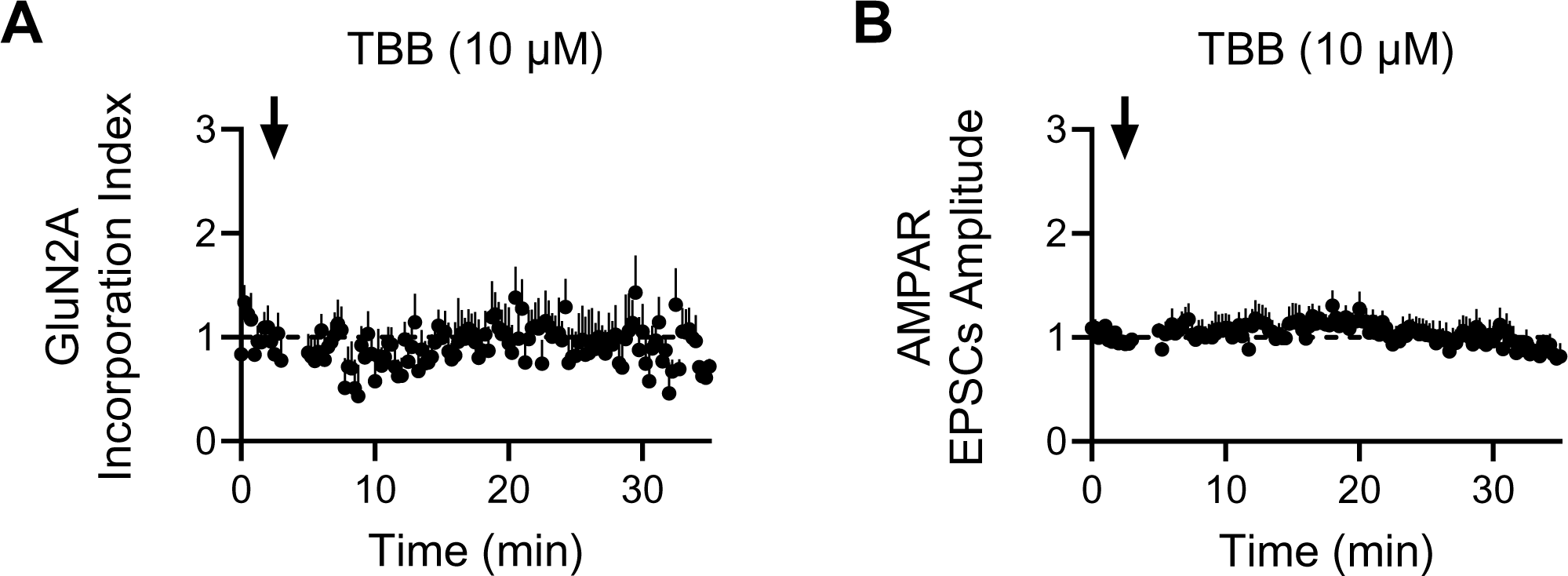
Inhibition of Casein kinase II and its effect on GluN2A synaptic incorporation. **A.** Amplitude of the current carried by GluN2A-containing recombinant NMDARs over time in CA1 neurons co-transfected with GFP-tagged NMDAR subunits GluN2A and GluN1 N598R. Slices were incubated with TBB, a Casein Kinase II inhibitor, 2 hours before the experiment (n=8). EPSCs were recorded at -70 mV and the GluN2A current measured 150 ms after the stimulus artifact. The arrow indicates the induction of synaptic potentiation through a pairing protocol, which consists of 2-minute, 3 Hz stimulation while holding the cell at 0 mV. **B.** Peak amplitude of EPSCs recorded at -70 mV, representing the AMPAR component of the EPSC, in CA1 neurons coexpressing GFP-tagged NMDAR subunits GluN2A and GluN1 N598R, and treated with TBB for 2 hours before the experiment.

Together, these experiments indicate that the removal of GluN2B receptors in an activity-dependent manner is necessary to allow for the synaptic incorporation of GluN2A receptors. Additionally, they suggest that the potentiation of AMPAR-mediated synaptic transmission is linked to the synaptic incorporation of NR2A-containing NMDA receptors.

## DISCUSSION

The expression of glutamate receptors at synaptic sites follows a well-orchestrated choreography, characterized by specific temporal and regional patterns in the production of the various subunits that make up both AMPA-type and NMDA-type receptors. These different subunits endow the ion channels with distinct biophysical properties and determine their capacity to interact with specific postsynaptic scaffolding and signaling proteins (Dingledine et al., 1999; Traynelis et al., 2010; Hansen et al., 2021).

The precise modulation of synaptic currents and the interactions with postsynaptic proteins play a crucial role in regulating synaptic plasticity (Barria and Malinow, 2005; Zhou et al., 2007), which is essential for the proper selection of synaptic contacts and the development of neuronal circuits. Consequently, disruptions in the expression and synaptic incorporation of glutamate receptor subunits during early synaptogenesis can lead to aberrant neural wiring, potentially manifesting later in life as cognitive disorders such as Fragile X syndrome or schizophrenia (Banke and Barria, 2020; Banke et al., 2024).

Of particular importance is the switch in GluN2 subunits of synaptic NMDARs. Receptors containing either GluN2A or GluN2B follow different paths to reach synapses and are also removed in a different manner from them.

While GluN2B-containing receptors can reach synapses independently of synaptic activity, GluN2A-containing receptors require a certain level of synaptic activity for their incorporation into synapses (Barria and Malinow, 2002). GluN2B is often expressed extrasynaptically, where it has a high diffusion coefficient (Groc et al., 2006; Kellermayer et al., 2018), allowing it to be incorporated into synapses through lateral diffusion (Gambrill et al., 2011; McQuate and Barria, 2020). This same mechanism allows GluN2B to leave synapses, though it can also be internalized via clathrin-dependent endocytosis through a YEKL motif in its carboxy-terminal domain that interacts with the AP2 complex (Lavezzari et al., 2003; Lavezzari et al., 2004). Additionally, GluN2B has been found to increase spine and filopodia motility (Gambrill and Barria, 2011).

In contrast, GluN2A-containing receptors seem to be trafficked directly to dendritic spines—the primary sites of synaptic contact in most excitatory neurons. They tend to accumulate in these locations if the binding of glutamate to preexisting synaptic receptors is blocked, which prevents their synaptic incorporation (Barria and Malinow, 2002). Once at the synapse, GluN2A-containing receptors are generally more stable, and it has been proposed that they confer stability to synapses and spine structures (Foster et al., 2010; Gambrill and Barria, 2011).

Changes in the decay kinetics of isolated NMDAR-mediated EPSCs, along with sensitivity to ifenprodil, a specific non-competitive antagonist of GluN2B, clearly indicates that as synapses mature, the ratio of GluN2B to GluN2A decreases. However, it remains unclear whether this involves a direct one-to-one exchange of GluN2B for GluN2A, or if GluN2B leaves synapses through simple diffusion or an activity-dependent mechanism involving receptor endocytosis.

Since synaptic incorporation of GluN2A requires glutamate binding to preexisting NMDARs, it has been proposed that clathrin-mediated removal of GluN2B creates a synaptic spot for GluN2A insertion. The observation that long-term potentiation at hippocampal glutamatergic synapses results in faster decay kinetics of NMDAR-mediated EPSCs without a change in their peak amplitude (Bellone and Nicoll, 2007) also supports the idea of a one-to-one exchange between GluN2B and GluN2A.

Our results directly tested this hypothesis by blocking the clathrin-dependent removal of GluN2B-containing receptors. By blocking the endocytosis of GluN2B-containing NMDARs through the expression of the YEKL peptide in CA1 neurons, and measuring the synaptic incorporation of recombinant GluN2A, which is not inhibited by Mg^2+^, we demonstrate that these two processes are directly linked, supporting the hypothesis that the removal of GluN2B is necessary for the incorporation of GluN2A at synapses.

By employing a protocol that enables the expression of Mg^2+^-insensitive GluN2A-containing receptors while preventing their synaptic incorporation (Barria and Malinow, 2002), we could also examine specific conditions necessary for GluN2A’s synaptic incorporation in real time.

Understanding the conditions required for GluN2A synaptic incorporation is crucial, as it has been observed that in hippocampal slices treated with tetrodotoxin (TTX) for 2-3 days, which blocks synaptic activity but not spontaneous neurotransmitter release, GluN2A is still incorporated into synapses. This suggests a mechanism that removes existing NMDARs without activating their ion channels and, therefore, operates in a Ca^2+^-independent manner (Barria and Malinow, 2002).

Our results demonstrate that the rapid synaptic incorporation of GluN2A induced by a commonly used long-term potentiation (LTP) protocol requires postsynaptic Ca^2+^, indicating that mere glutamate release is not enough to trigger the switch in subunit composition.

It is possible that stimulation of preexisting NMDARs by spontaneous release of glutamate during the 2-3 days of TTX incubation still can produce small Ca^2+^ transients necessary for the incorporation of GluN2A, or that a slower activity-independent synaptic incorporation of GluN2A exist.

Notably, blocking GluN2B internalization—whether by expressing the YEKL peptide or using rapid blockade with Ant-YEKL peptide or TBB—not only prevents the incorporation of GluN2A into synapses, but also inhibits the potentiation of synaptic currents mediated by AMPARs. This suggests that long-term potentiation (LTP) not only boosts AMPAR trafficking (Kessels and Malinow, 2009) but also incorporates GluN2A. Such a process could promote synaptic structure stability, as studies suggest that dendritic spines containing GluN2A are more stable than those with GluN2B (Gambrill and Barria, 2011). Whether GluN2A and AMPAR are incorporated from the same intracellular compartments remains a topic for future investigation.

These findings shed light on the molecular mechanisms driving the switch from GluN2B to GluN2A in NMDA receptors. They also reveal how functional interactions between various glutamate receptors in the brain may regulate the levels of AMPARs and contribute to synaptic potentiation and stabilization.

## Acknowledgments

This work was supported by the National Science Foundation (NSF) grant NSF-IOS-2224262 (A.B.).

