## Supplemental Figure for "ACTIVITY-DEPENDENT INTERNALIZATION OF GLUN2B-CONTAINING NMDARS IS REQUIRED FOR SYNAPTIC INCORPORATION OF GLUN2A AND SYNAPTIC PLASTICITY"

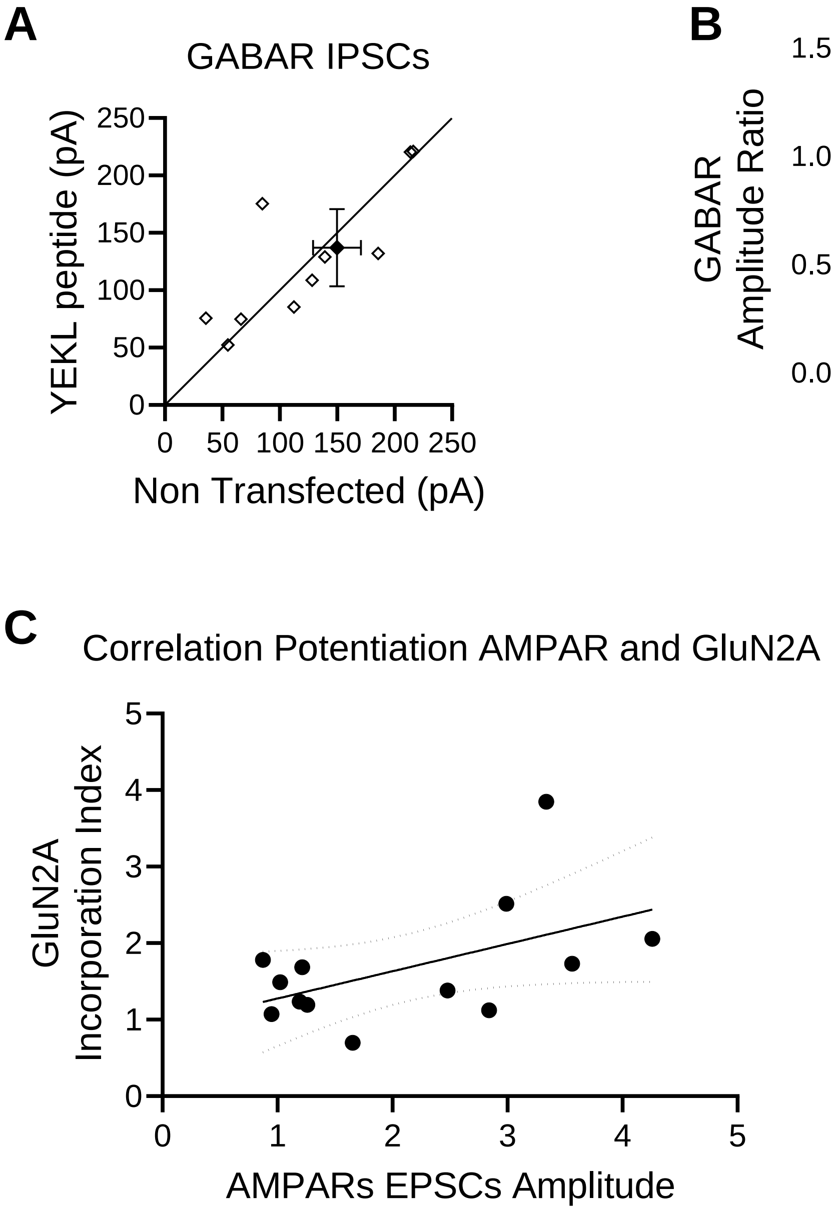


Supplemental Figure

**Supplemental Figure**

**A.** Paired recordings of neurons transfected with YEKL peptide and control non-transfected (NT) neurons. Evoked Inhibitory Postsynaptic Currents (IPSCs) were recorded in the presence of 100 μM APV and 2 μM NBQX (n= 11 pairs). Optically tagged YEKL peptide was transfected using biolistic and expressed for 2-3 days. Black dot is average ± s.e.m.

**B.** Peak amplitude of IPSCs from paired recordings normalized to the amplitude of the control non-transfected (NT) neuron. Statistical significance was determined using Wilcoxon matched pairs test (p=0.6377).

**C.** Correlation between AMPAR potentiation after pairing protocol and synaptic incorporation of GluN2Acontaining recombinant NMDARs. Line is linear regression fit with 95% confidence interval.
